# High levels of origin licensing during *Xenopus* cleavage divisions ensures complete and timely genome duplication

**DOI:** 10.1101/2021.10.12.464070

**Authors:** Peter J. Gillespie, Jolanta Kisielewska, Mohammed Al Mamun, Guennadi Khoudoli, Kevin Creavin, Alan J. Score, J. Julian Blow

## Abstract

Cells face several challenges to completing genome duplication. One challenge is the irreversible stalling of converging replication forks (‘double fork stalls’). Cell types that cannot delay mitotic entry must also ensure that no replication origins are too far apart (the ‘random gap problem’). We show how these challenges can be met in early *Xenopus* embryos by the very abundant licensing of replication origins: one MCM2-7 double hexamer every ∼250 bp. Licensing does not change nucleosome spacing, consistent with MCM2-7 being assembled onto inter-nucleosomal linker DNA. We show that later embryonic development can occur successfully with a per-cell cycle failure rate of <0.2% in early embryos. The high density of licensed origins in the early embryo reduces cell cycle failures from random gaps and from double fork stalls to levels compatible with subsequent development, suggesting that *Xenopus* early embryonic cells can ensure complete genome duplication without requiring unconventional replication mechanisms.

## Introduction

The need to achieve complete genome duplication during each eukaryotic cell division cycle presents a number of major challenges. Replication forks may irreversibly stall on encountering DNA breaks, modified bases or tightly bound proteins; in addition replication origins may be too few in number, may be inappropriately located on the genome or may not be activated in a timely manner. These problems are exacerbated by the fact that the number and position of potentially active origins must be specified before cells enter S phase. This is because the loading of MCM2-7 proteins onto replication origins – origin licensing – occurs only during anaphase and G1 as part of an essential mechanism to prevent the re-replication of segments of DNA^1-3^. Whatever problems are encountered during S phase, the only replication origins that can be used are the ones that had already been licensed at S phase entry.

Because replication is initiated bidirectionally from origins, an isolated fork stall by itself does not create an insurmountable problem because the intervening DNA can be replicated by the fork emanating from the downstream origin (except at the most telomeric origin)^4^. However, if both of the converging forks irreversibly stall and there is no licensed origin between them, this creates a Double Fork Stall (DFS) which cannot be resolved by standard mechanisms (Fig 1a). We have previously shown that in organisms with smaller genome sizes such as yeasts (genome size ∼10 Mbp) this problem can be solved by the appropriate distribution of replication origins, spaced on average 20 kb apart^4^. However, with similar average inter-origin spacing the much larger size of metazoan genomes will experience a DFS in a high proportion of cell cycles, necessitating additional mechanisms for completing replication^5-7^. We and others have provided evidence that metazoans have evolved mechanisms by which the complete replication of their large genomes can be delayed until mitosis and the early parts of the following cell cycle^5,6,8-10^.

**Figure 1.**
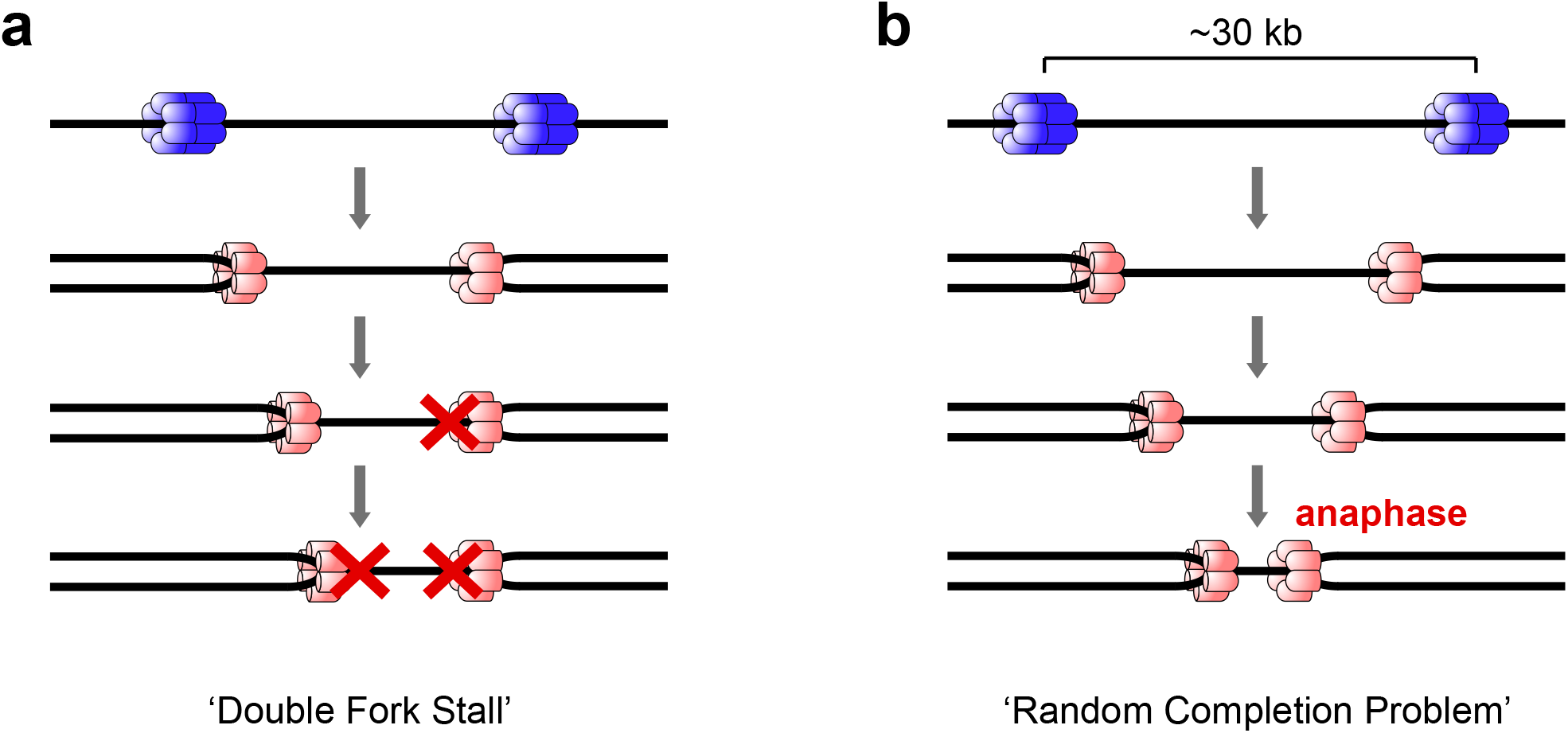
Potential problems in Xenopus embryos that result in incomplete genome duplication. Cartoon of two replication origins in the body of a chromosome. Double-stranded DNA is denoted as a single black line. Prior to S phase entry, origins are licensed by binding a double hexamer of MCM2-7 proteins (blue); as an origin fires, both MCM2-7 single hexamers are converted into an active CMG helicase (pink). A. If converging replication forks both stall (red cross) and there is no dormant origin between the stalled forks then the resultant ‘double fork stall’ leaves the intervening DNA unreplicated during S phase. B. ‘Random Completion Problem’. In the early *Xenopus laevis* embryo entry into S phase is not delayed if replication is incomplete. Since there is only ∼25 mins for S phase and together with a replication fork speed of ∼10 nt sec^-1^, this means that if randomly distributed replication origins are greater than ∼30 kb apart then the intervening DNA cannot be replicated before chromatids are pulled apart in anaphase.

Since this mechanism for resolving unreplicated DNA may be time-consuming and have a limited capacity^6^, this may cause problems for cells with extremely short cell cycles, such as those in some early embryos (for example *Xenopus* and *Drosophila*). In this paper we consider how cells in the early *Xenopus* embryo might ensure complete genome replication. After fertilisation, *Xenopus* embryos undergo 12 rapid and synchronous cell division cycles in ∼7 hours before progression through the Mid-Blastula Transition (MBT) after which cells undergo slower asynchronous divisions more typical of somatic cells^11,12^. During these pre-MBT cleavage divisions, there is no cell growth, there is virtually no transcription and the early embryo essentially oscillates between S phase and mitosis. Importantly, cell cycle checkpoints are absent or weak prior to the MBT, so that even if S phase is incomplete, the cell cycle continues unperturbed^13-15^. Another noteworthy feature of these cleavage divisions is that any DNA introduced into the egg can be incorporated into a nucleus and replicated, with replication initiation taking place without a requirement for any special DNA sequence^16-20^. The basic nuclear events of the early embryonic cell cycle can be recapitulated in vitro by extracts prepared from Xenopus eggs^21-24^.

It has previously been reported that active replication origins in Xenopus egg extracts are spaced on average ∼10 kb apart^20,25^. However, only a fraction of all licensed origins are activated in any given S phase, and there are at least 10 times more origins licensed in Xenopus eggs and egg extracts than undergo replication initiation^26-29^. In particular, when both replication fork progression and checkpoint kinases are inhibited, replication origins are activated at very high density, in excess of 1 per kilobase^29^.

With a system in place that ensures that all licensed origins can potentially fire^30^ protection against DFSs is determined by the number and distribution of licensed origins irrespective of the total number that actually fire^31,32^. One of the questions we pose here is whether the number and distribution of licensed origins in the Xenopus early embryo is sufficient to reduce the probability of DFSs to acceptable levels prior to the commencement of the more normal divisions after the MBT.

A second problem arises because of the rigid timing of cell divisions prior to the MBT. With a fixed cell cycle duration of 30 mins that is insensitive to incomplete replication^13,14^, a fork rate of ∼600 bp/min^19^ and a minimum time of 5 mins for mitosis, then if replication origins are more than ∼30 kb apart there will not be enough time for the DNA between the two origins to be completely replicated before chromosomes are segregated in anaphase. This is the ‘random completion problem’^18,25,33^: with origins placed at random with respect to DNA sequence, how do cells ensure that no licensed origins are more than ∼30 kb apart (Fig 1b)?

Here we provide evidence that approximately four MCM2-7 double hexamers are assembled on chromatin in Xenopus egg extracts per kb of chromatin. This occurs without significant disruption to the normal chromatin structure, consistent with the idea that MCM2-7 proteins are loaded onto linker DNA between nucleosomes. This high density of licensed origins on DNA can explain why replication initiation occurs apparently without regard to DNA sequence in the early embryo. We then show that approximately normal embryonic development can occur with a cell cycle failure rate of ∼0.2% in pre-MBT divisions. Mathematical modelling suggests that the density of MCM2-7 we report is sufficient to bring the probability of DFSs and large unlicensed gaps of >30 kb to levels low enough to support embryo survival until the MBT.

## Results and Discussion

### Capacity for licensing in Xenopus egg extract

The licensing of replication origins requires four essential factors: ORC, Cdc6, Cdt1 and MCM2-7^34-36^. ORC, Cdc6 and MCM2-7 are all ATPases, and their hydrolysis of ATP can change the way they interact with DNA^34,37-40^. We examined the effect of using apyrase to remove ATP from Xenopus egg extract and replacing it with a non-hydrolysable ATP analogue, ATP-γ-S. As a control, we also supplemented the egg extract with geminin, a natural inhibitor of licensing^41,42^. Consistent with previous reports, when licensing was blocked with either geminin^28,43^ or ATP-γ-S^34^ ORC and Cdc6 were stabilised on chromatin (Supplementary Figure S1). We also investigated the consequence of adding ATP to the buffer in which chromatin is isolated. When ATP was included in the chromatin isolation buffer under conditions where licensing could occur, almost no ORC or Cdc6 were isolated on chromatin (Supplementary Figure S1), consistent with the idea that ATP hydrolysis by these proteins releases them from DNA^37^. However, when licensing was blocked with either geminin or ATP-γ-S, isolated chromatin still contained maximal levels of ORC and Cdc6 even when isolated in buffer supplemented with ATP. We therefore decided that to monitor the recruitment of ORC and Cdc6 onto chromatin, we would replace ATP in the extract with ATP-γ-S.

In Xenopus egg extracts, ORC and Cdc6 are recruited rapidly to DNA, reaching a maximum within 1-2 minutes, but MCM2-7 loading (origin licensing) proceeds at a slightly slower pace reaching a maximum in 10 - 20 minutes^28^. We examined the saturation of ORC and Cdc6 on chromatin when licensing was blocked with ATP-γ-S. Equal quantities of DNA were incubated for 20 minutes with increasing volumes of egg extract; chromatin was then isolated and immunoblotted for ORC and Cdc6. Fig 2 shows that the amount of Orc1 loaded onto chromatin increased with the amount of extract added, until a plateau was reached at ∼30 μl extract per μg DNA. At this plateau, the amount of Orc1 on DNA was ∼15 times more than is present in 0.5 μl extract; since we know that Orc1 is present in extract at ∼50 nM^44^ we calculated that this represents ∼1.3 molecules of Orc1 bound per kb of DNA (see Materials and Methods for details of the calculation). Cdc6 did not show such a clear plateau as Orc1, but reached a maximum of ∼5 times the amount in 0.5 μl extract. The concentration of Cdc6 in egg extract is ∼80 nM^45^, allowing us to estimate that this represents ∼0.6 molecules of Cdc6 bound per kb of DNA (see Materials and Methods for details of the calculation).

**Figure 2.**
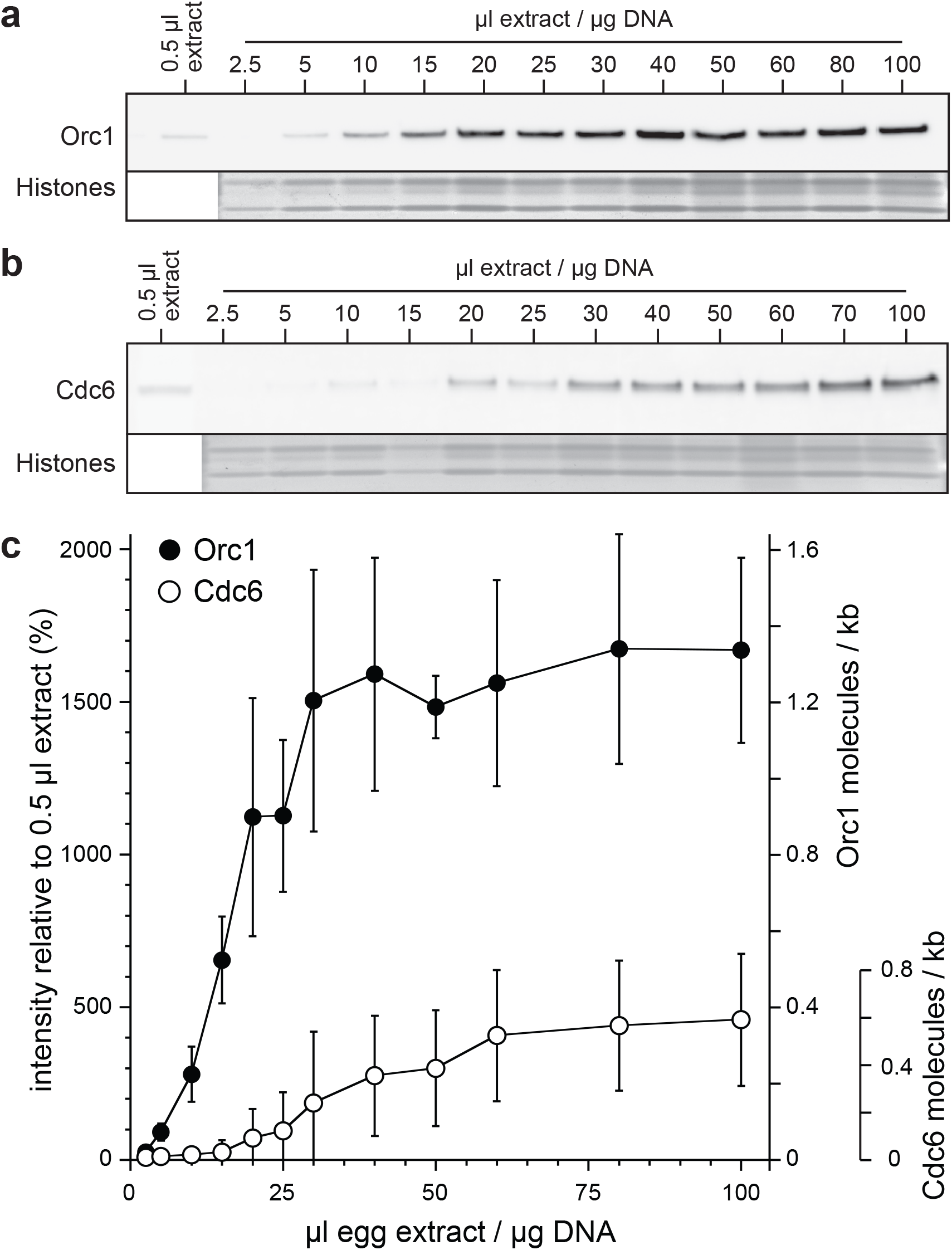
Quantification of Orc1 and Cdc6 chromatin saturation when replication licensing is inhibited by ATP-γ-S. **A, B**. Immunoblots showing the association of Orc1 (A) and Cdc6 (B) with *Xenopus laevis* sperm chromatin incubated for 20 mins in increasing volumes of apyrase-treated interphase *Xenopus* egg extract supplemented with 2.5 mM Mg•ATP-γ-S to inhibit licensing. C. The intensity of Orc1 and Cdc6 on chromatin was quantified and expressed relative to 0.5 μl of egg extract and the saturation per kb of DNA. n = 3 and error bars represent SEM.

In Xenopus, Cdc6 binding is dependent on ORC^45^, and the data in Fig 2 therefore suggests that at saturation, only about half of the ORC bound to DNA is associated with Cdc6. Since ORC and Cdc6 act catalytically and can cycle on and off DNA, each potentially loading multiple MCM2-7 hexamers onto DNA, we imagine that the density of MCM2-7 on DNA at saturation will be even higher than the density of ORC and Cdc6.

In order to estimate the density of MCM2-7 on DNA at saturation, we used quantitative western blotting to obtain an estimate for the concentrations of MCM3 and MCM6 in the egg extract. We used serial dilutions of the recombinant polypeptides used to make the antibodies for the MCM3 and MCM6 antibodies and then compared the intensity of the signal against serial dilutions of egg extract. MCM2-7 are very abundant proteins in Xenopus eggs, and we noted that in order to give a signal that was linearly proportional to the amount of protein on immunoblots, very small amounts of egg extract needed to be run on each gel lane (no more than 100 nl). Representative blots are shown in Supplementary Figure S2. These experiments provided estimates of 18 μM for MCM3 and 16 μM for MCM6 in Xenopus egg extracts. These values are higher than previous estimates^26^ which we believe is due to more sensitive detection of MCM2-7 in our current study due to the significantly lower protein concentrations and volumes used. Furthermore, our previous study used X-ray film, which is inherently non-linear, whilst in this study we either used fluorescent antibodies with images captured with a LiCor Odyssey or enhanced chemiluminescence with images captured on a CCD camera, both of which facilitate linear quantitative output over at least a 100-fold range.

The values that we calculate for MCM3 and MCM6 protein concentration in egg extract are greater than those derived from two studies in which protein concentration in Xenopus egg extract was determined by quantitative proteomics: MCM3 0.5-0.7 μM and MCM6 0.3-0.6 μM^46,47^. It should be noted that the methods of extract preparation and the buffers used in these two studies are significantly different from the method that we use. In the current study we used previously optimised extract methods for maximising the efficiency of replication licensing activity: including lengthy ultracentrifugation steps in which only ∼1/8 of the starting volume of eggs is recovered as usable cytoplasm and buffer conditions that reduce the concentration of Cl^-^ ions (replacing them with HEPES and phosphate anions) because of the instability of the MCM2-7 complex in the presence of small anions. It should therefore be noted that neither proteomic study uses extraction conditions optimal for the maintenance of licensing activity or MCM2-7 complex integrity; in particular the study in which Tris is used for extraction^46^ the overall recovery of MCM2-7 is lower than that in which HEPES alone was used^47^ and specifically, the recovery of MCMs 4, 6 and 7 are significantly less than that of MCMs 2, 3 and 5; this suggests that the MCM2-7 complex has fragmented and at least some of the MCM4-6-7 subcomplex has been lost by precipitation on extraction^48^.

### Abundance of MCM2-7 on embryonic chromatin

To determine the amount of MCM2-7 loaded onto DNA, we performed experiments in Xenopus extract without added ATP-γ-S or geminin. Under normal circumstances licensing occurs from the time when geminin is inactivated at the start of anaphase^42,49^ until nuclear import of geminin reactivates it as a licensing inhibitor which typically takes 20-30 minutes after DNA is added to egg extracts^50-52^. Close to the MBT, replication-dependent degradation of Cdt1 further enhances the block to re-replication^52-54^. In order to determine whether the reactivation of geminin or degradation of Cdt1 limits the total amount of MCM2-7 loaded onto DNA, we isolated chromatin from extracts that had optionally been supplemented with Wheat Germ Agglutinin (WGA) to block nuclear pore function^55^. Despite the WGA completely blocking nuclear assembly (Supplementary Figure S3), it caused no significant reduction in the total amount of MCM3 or MCM6 loaded onto chromatin after 20 mins (Fig 3a and 3b). We therefore conclude that in egg extracts chromatin becomes saturated with MCM2-7 before nuclear assembly shuts down the licensing system.

**Figure 3.**
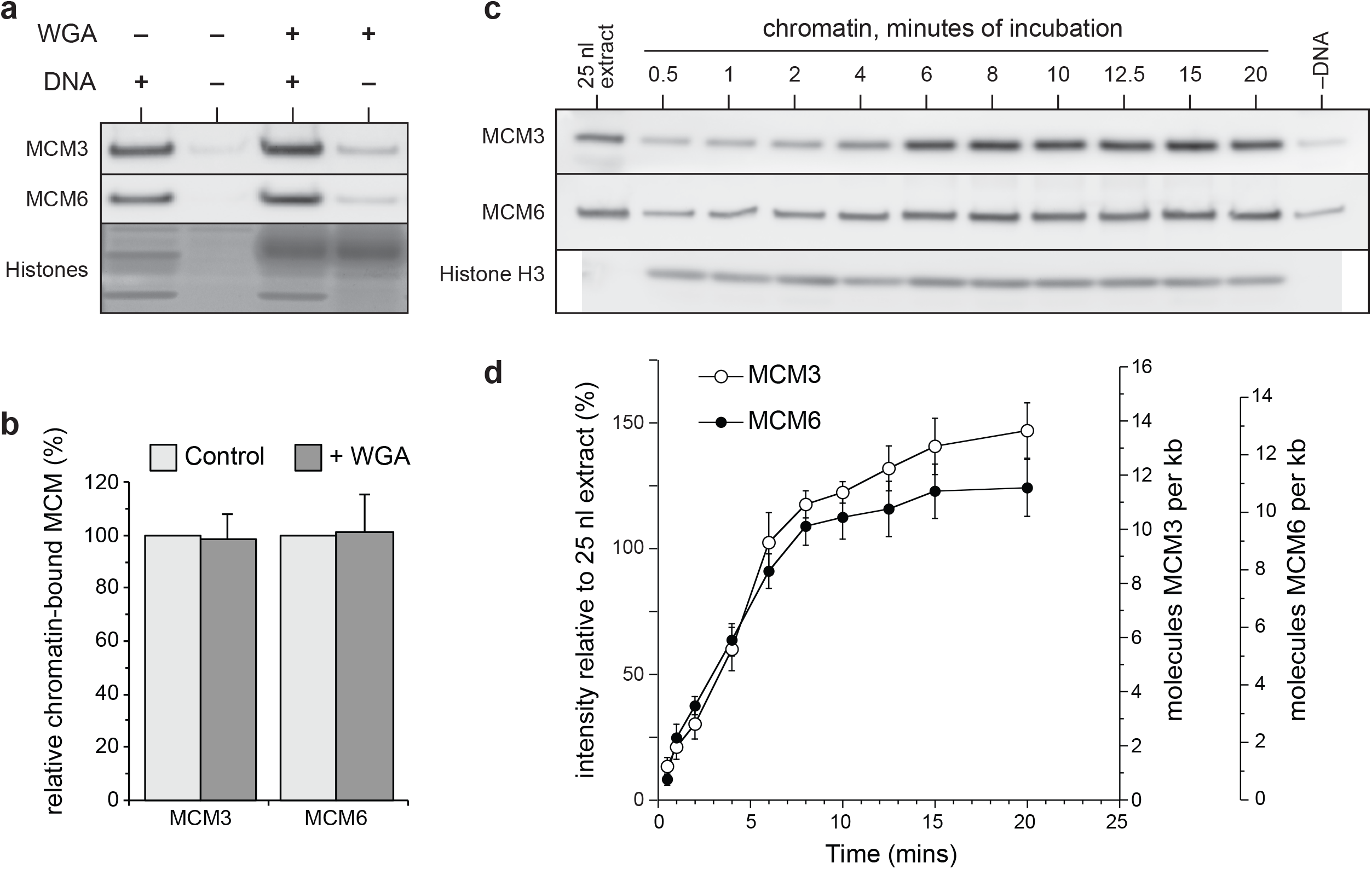
The kinetics of replication licensing in vitro: immunoblotting. Kinetics of MCM3 and MCM6 association with sperm chromatin during the licensing period in interphase *Xenopus* egg extract, prior to nuclear envelope assembly. A. Quantitative immunoblot of MCM3 and MCM6 recovered from interphase egg extract optionally supplemented with sperm chromatin (DNA) and/or wheat germ agglutinin (WGA). The lower portion of the gel was stained with Coomassie Blue to visualise histones. B. The mean intensity of chromatin-bound MCM3 and MCM6 plus or minus (WGA) from (A) after subtraction of signal in lanes with no added chromatin; error bars represent SEM of 3 experiments. C. Quantitative immunoblot of MCM3, MCM6 and histone H3 association with sperm chromatin incubated for 20 mins in *Xenopus* interphase egg extract treated with wheat germ agglutinin. At the indicated times 100 nM of geminin^DEL^ was added to individual samples to arrest the licensing reaction. -DNA shows the background signal when no sperm was added. D. The mean intensity of chromatin bound MCM3 and MCM6 in (C) expressed relative to 25 nl egg extract and the saturation per kb of DNA. The values shown have had the signal in samples with no added chromatin subtracted from them; error bars represent SEM of 3 experiments. The number of molecules of MCM3 and MCM6 loaded per kb of DNA was calculated assuming undiluted extract contains 18 μM MCM3 and 16 μM MCM6.

We then performed a time course in which licensing was allowed to occur for different periods of time after which constitutively active geminin (gemininDEL)^41^ was added to stop further licensing. To prevent reactivation of endogenous geminin following nuclear import, extract was also supplemented from the start with WGA. Chromatin was isolated at 20 mins - the earliest time at which nuclear assembly might be expected to be completed - and blotted for MCM3 and MCM6 (Fig 3c). The rate of MCM3 and MCM6 loading started off high but then appeared to reach plateau values after ∼10 minutes. We quantified the amount of MCM3 and MCM6 loaded relative to the extract loading control (25 nl) and then used the previously-derived concentrations of these proteins to estimate the total amount of protein loaded onto chromatin (Fig 3d). This indicated that up to 10 - 14 molecules of MCM3 and MCM6 were loaded per kb of DNA. Since these proteins are loaded onto DNA as double hexamers of MCM2-7^35,36,56^ this suggests that up to 5 - 7 MCM2-7 double hexamers were loaded onto each kb of DNA in these experiments. Under these conditions, significantly less than half of the total stockpile of Orc1, Cdc6, Cdt1 and MCM3 in the extract had been loaded onto the DNA assembly (Supplementary Figure S4).

We were surprised by this very high density of MCM2-7 on chromatin indicated by immunoblotting and therefore set out to measure it by a different means. When MCM2-7 hexamers encircle DNA to license the DNA for replication, the hexamers are relatively stable to high salt concentrations. We therefore isolated the maximally licensed chromatin from Xenopus egg extract and carefully washed it with salt to remove weakly bound protein. As controls we isolated chromatin from extract treated with geminin to block licensing (‘geminin chromatin’) or isolated ‘mock chromatin’ from extract to which no DNA had been added. The chromatin samples were then separated on protein gels and visualised by SYPRO Ruby staining, as shown in Fig 4a^57^. Protein bands at the expected position of MCM2-7 and the four core histones were clearly visible; the identity of these proteins was confirmed by eluting the proteins from gel slices and subjecting them to mass spectrometry. The bands corresponding to MCM2-7 were absent from chromatin isolated from the geminin-treated extract and were also absent from the ‘mock chromatin’. The bands corresponding to core histones were also absent from the ‘mock chromatin’. We also noted several bands in the geminin sample that were absent from the licensed chromatin. Previous reports have shown that ORC and Cdc6 bind more tightly to chromatin when licensing is blocked^28,43^, consistent with the results shown in Supplementary Fig S1. We therefore cut out some of the bands exclusive to the geminin chromatin and analysed them by mass spectrometry. As indicated in Figs 4a and 4b, we could identify bands corresponding to Orc1-3 in the unlicensed chromatin prepared from geminin-treated extract as well as strong bands corresponding to TopoII and the chromatin remodelling enzymes ISWI, Baz1a and Baz1b.

**Figure 4.**
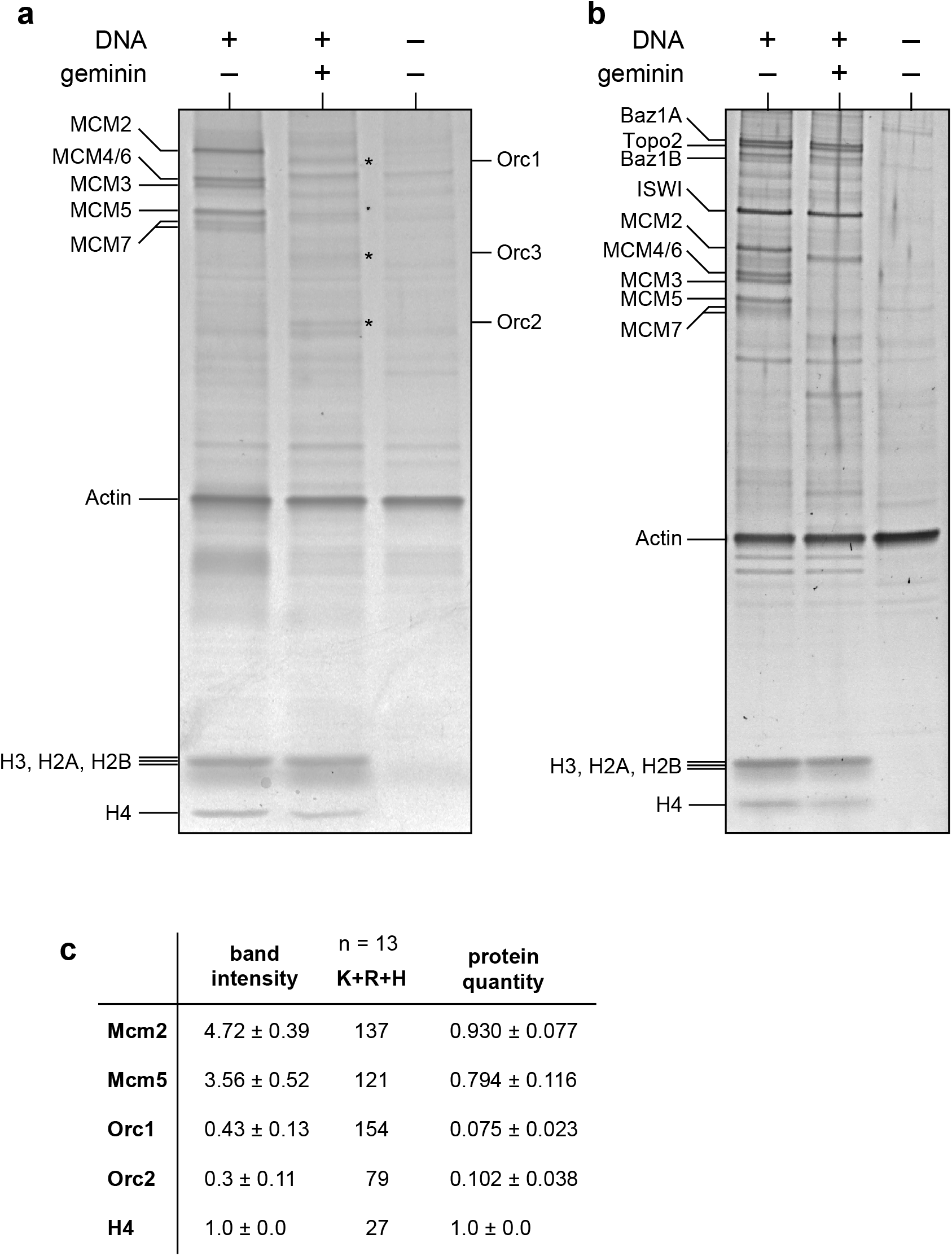
Quantification of replication licensing in vitro using SYPRO-Ruby staining of isolated chromatin. Quantification of protein recovered from *Xenopus* egg extract incubated for 20 mins and optionally supplemented with sperm chromatin and/or the licensing inhibitor geminin. **A, B**. SYPRO Ruby stained SDS-PAGE gels of recovered proteins. MCM2-7, Orc1-3 (asterisked to aid visibility), ISWI and ISWI subunits Baz1A and Baz1B, Topo II and the histones are indicated. Contaminating actin which is recovered in the absence of added chromatin is indicated. C. The abundance of MCM2, MCM5, Orc1 and Orc2 on chromatin was quantified relative to Histone H4. The intensity of each SYPRO Ruby stained protein band was determined together with that of the equivalent position in the ‘no chromatin added control’ which served as background and was subtracted from the value determined for each protein. The total arginine (R), lysine (K) and histidine (H) content of each protein was calculated and then used to the quantify the amount of each protein relative to Histone. n = 13 and SD is indicated.

We next quantified the intensity of the protein bands corresponding to MCM2, MCM5, Orc1, Orc2 and histone H4. We chose these 5 proteins as they can be identified as clear single bands on the gel. The intensity of these proteins is tabulated in Figure 4c, expressed relative to the intensity of histone H4. SYPRO Ruby, which was used to stain the protein bands, binds primarily to lysine (K), arginine (R) and histidine (H)^58^. The abundance of these amino acids in the 5 proteins is also shown in Fig 4c and can be used to determine the relative concentrations of different proteins^58,59^. We therefore used the H+K+R abundance in each protein to estimate the molar abundance relative to histone H4 (Fig 4c, ‘protein quantity’). This indicated that the amount of MCM2 and MCM5 on chromatin was 79-93% that of histone H4. Since both proteins are present in dimeric complexes (two H4 per nucleosome and two copies of each MCM protein in a double hexamer) this means that there is just under one MCM2-7 double hexamer per nucleosome. The nucleosome spacing in Xenopus early embryos is 180 - 200 bp^60-62^ meaning that there are 5 - 5.5 nucleosomes per kb. If MCM2-7 is 85% as abundant on chromatin as histone H4, this means that there will be 4.2 - 4.7 double hexamers per kilobase, a figure that is in good agreement to that derived from quantitative immunoblotting (Fig 3). Using the same approach, we estimated the quantity of Orc1 to be 7.5% of H4 and Orc2 to be 10% of H4 (Fig 4c). This suggests that ORC is present on unlicensed chromatin at ∼1 copy per kb, similar to the figure derived from quantitative immunoblotting (Fig 3). Once licensing has occurred, ORC and Cdc6 binding reduces significantly (Supplementary Figure S1)^28,43^.

The DNA template used for these studies is demembranated Xenopus sperm, the natural substrate for replication in egg cytoplasm. Xenopus sperm chromatin contains approximately normal levels of histones H3 and H4, but most H2A and H2B are replaced by protamines^61^. When added to egg extract, the protamines are rapidly exchanged for H2A and H2B by the histone chaperones nucleoplasmin and N1/N2, causing the compacted sperm chromatin to decondense; this decondensation is required to support the licensing of the chromatin^61,63-65^. We considered whether the unusual chromatin structure of sperm chromatin might account for the very high loading of MCM2-7. We therefore fully replicated sperm chromatin in Xenopus egg extract (at which time all nucleosomes would consist of the canonical H3-H4-H2A-H2B structure), drove the extract into mitosis to allow relicensing of DNA, released the mitotic extract back into interphase and then isolated the chromatin. After salt-washing and separation of proteins as before, the relative quantity of MCM2 and MCM5 remained at 87% and 72% compared to H4 (Supplementary Figure S5). We therefore conclude that the extremely dense loading of MCM2-7 onto chromatin, at ∼4 copies per kb, is a general feature of early embryonic chromatin. This very high density of potential replication origins provides an explanation of why virtually any DNA sequence can act as a replication origin in the Xenopus early embryo^16-19^.

With this very large amount of MCM2-7 on chromatin, we wondered whether it would disturb nucleosomal structure. We therefore prepared chromatin from extract prior to S phase plus or minus added geminin and also replicated G2 chromatin (lacking MCM2-7 following the completion of genome duplication). Chromatin was digested with micrococcal nuclease and the DNA was separated on agarose gels. Representative examples are shown in Figure 5a. In all samples, the 146 bp fragment representing the core nucleosomal DNA is clearly evident, plus a ladder of fragments with a spacing of ∼200 bp. Although there are subtle difference between the licensed, unlicensed and G2 chromatin that can be observed, the overall patterns are remarkably similar.

**Figure 5.**
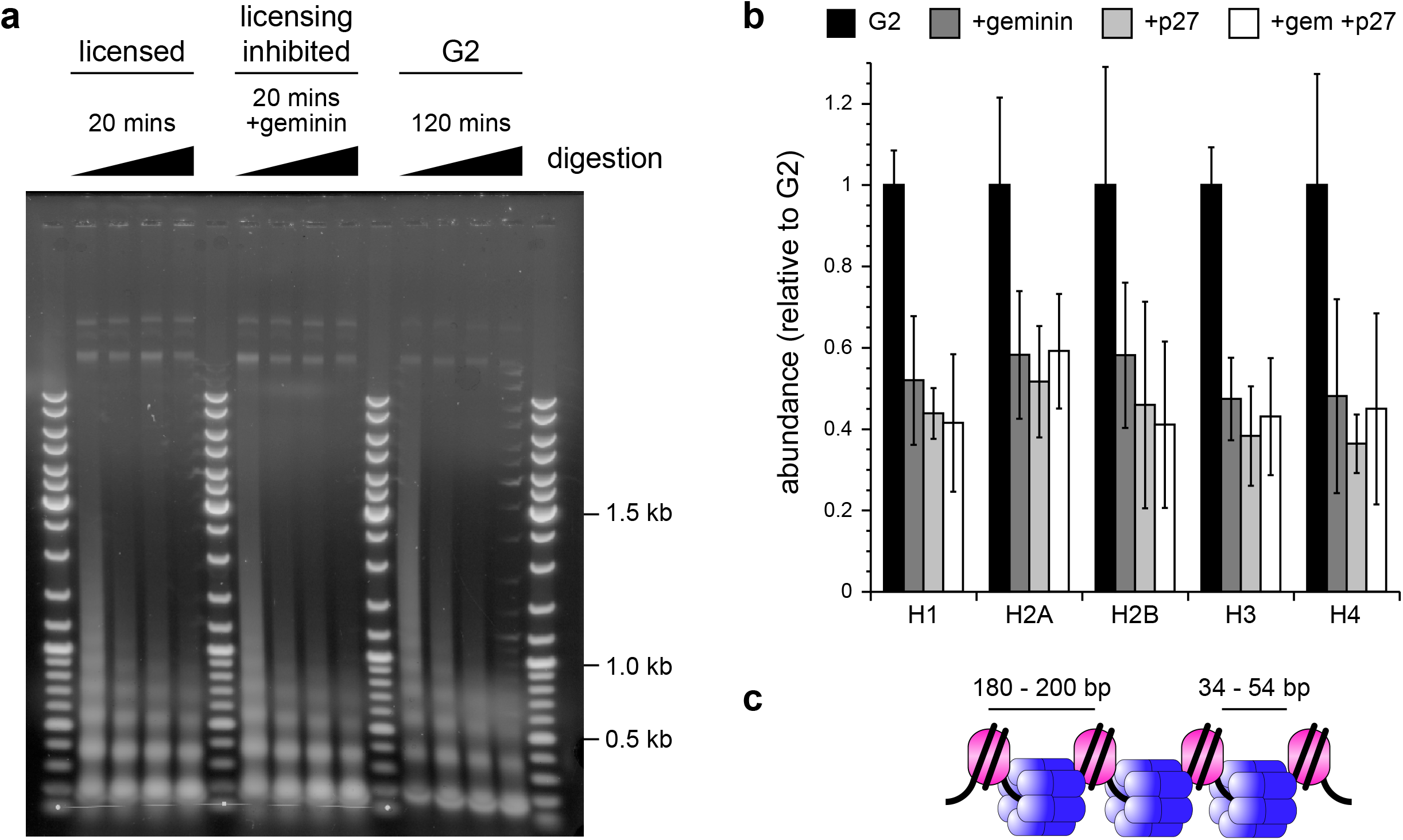
Characterisation of chromatin in egg extract during replication licensing. **A.** Micrococcal (MNase) digestion patterns of chromatin formed following incubation of *Xenopus laevis* sperm chromatin in interphase *Xenopus* egg extract for 20 mins (licensed) or 20 mins plus geminin (unlicensed) or for 120 mins (G2). DNA markers flanking each treatment are a 100 bp ladder. **B**. The extent of histone abundance on chromatin following incubation for 120 mins in egg extract optionally supplemented with geminin^DEL^ or p27^KIP1^ as determined by LC-MS. The intensity of each histone type was collated and expressed as a percentage of that present in the replicated control sample; error bars represent SEM of 3 experiments. **C**. Model depicting a possible alternating arrangement of double hexameric MCM2-7 and nucleosomes. Inter-nucleosomal distance and linker DNA length are indicated.

We next used mass spectrometry to examine the proteins associated with chromatin under different conditions, using a label-free approach to estimate protein abundance^66^. Sperm chromatin was incubated for 120 mins in extract optionally supplemented with geminin (to block licensing) and/or p27^Kip1^ (to inhibit S phase CDKs and therefore block DNA replication). To simplify the analysis, we grouped the histone variants into their major subgroups (H1, H2A, H2B, H3 and H4), and show their aggregated abundances in Figure 5b. All abundances were normalised to the value from untreated G2 chromatin. When replication was blocked with either geminin or p27^Kip1^ the quantity of histones was approximately halved reflecting that the samples contained half the amount of DNA. Compared to the p27^Kip1^-treated sample, the geminin-treated sample had slightly higher levels of all core histone proteins but the absolute change was only around 10% and was well within the error range of the experiment. We conclude therefore that although there may be minor changes to nucleosomes when licensing occurs, in terms of both nucleosome spacing and nucleosome content, any effects are relatively small. There is no evidence that the presence of MCM2-7 on the chromatin makes a large change to chromatin structure.

If the normal chromatin structure is maintained when MCM2-7 is loaded onto DNA, and if there is almost one MCM2-7 double hexamer per nucleosome, one possibility is that the MCM2-7 double hexamers are loaded onto the linker DNA between the nucleosome cores. MCM2-7 double hexamers have a footprint on DNA of ∼60 bp^67,68^. The nucleosome spacing in Xenopus early embryos is 180 - 200 bp and with 146 bp of DNA wrapped around the nucleosome, that leaves 34-54 bp of linker DNA to accommodate each MCM2-7 double hexamer (Fig 5c).

### Consequences of MCM2-7 abundance on replication completion

Figure 1 above shows the risks to the successful completion of genome duplication in the rapid cycles of the early *Xenopus* embryo caused by DFSs and by the ‘random completion problem’. Having shown in Figures 2-5 above the abundance of licensed origins in the early embryo, we next wanted to explore the consequences of this for successful genome duplication during all of the cleavage cycles prior to the MBT. Since no biological process can be consistently 100% accurate, we started by determining the Xenopus embryos’ ability to tolerate loss of cells before the MBT. The MBT occurs at 6 - 7 hr after 12 rapid and synchronous cell division cycles. To mimic occasional cell cycle failure, we killed single cells in pre-MBT embryos by microinjecting them with DNase I at the 2, 4, 8, 16, 32, 64 and 128 cell stages. We then examined subsequent embryo development at 6 and 7 hr (approximately the MBT) and at 24 hr after fertilisation. Representative pictures are shown in Fig 6a and the number of surviving embryos are shown in Fig 6b. The destruction of single cells at any stage caused a reduction in embryo survival which is clearly seen 24 hr after fertilisation. However, destruction of single cells at the 64 and 128 cell stages was clearly less deleterious to survival than destruction of single cells at earlier stages and more than 80% of these embryos were viable at 24 hr. At this level of cell loss, embryos at the MBT contain 98-99% of the normal number of cells (4032 or 4064 cells rather than the 4096 expected if there are no errors).

**Figure 6.**
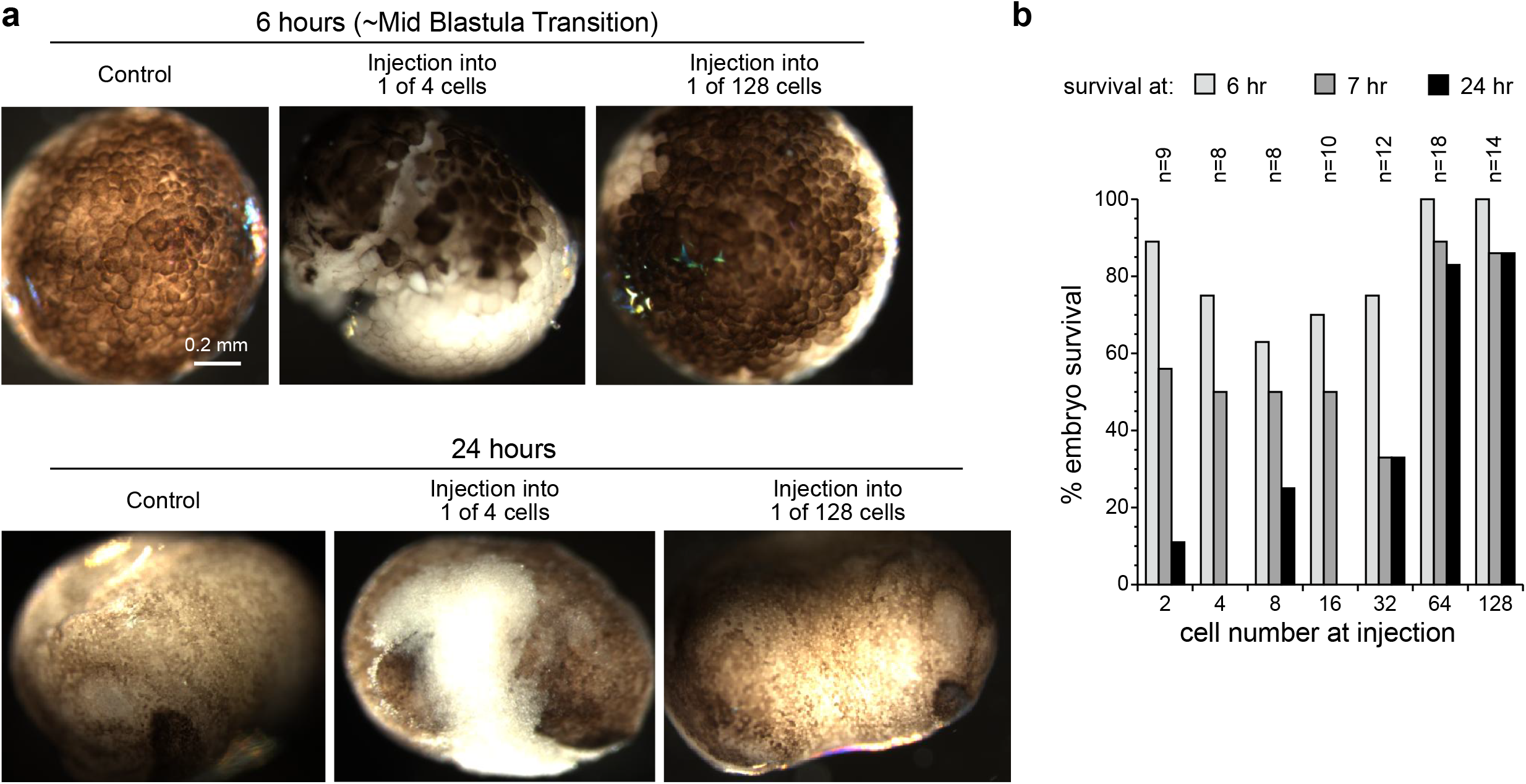
Xenopus embryo survival following loss of cell-lineage. Single cells in developing Xenopus eggs were microinjected with DNase to remove individual cell lineages at different cell numbers, prior to the mid-blastula transition. Development was followed over 24 hours. **A**. Representative photomicrographs depicting cell lineage removal at different cell number **B**. Embryo survival following microinjection was recorded at 6, 7 and 24 hours. The number of embryos injected for each measurement are indicated.

We next modelled how this degree of cell loss would correspond to a constant probability of failure in each cell cycle. The modelling was performed in two different ways. We first derived a simple mathematical equation where there is probability *p* of successful cell division. For each cell the expected average number o f c ells after a single round of division is therefore 2.*p*. After *n* divisions the expected number of cells *E(n)* = *2*^*n*^. *p*^*n*^.

The expected number of surviving cells after 12 cell cycles is shown in Figure 7a (black lines). This indicates that for embryos to reach the MBT with 4032 or more cells, the average failure rate should be less than ∼0.002. Because cell division is a series of discrete events not adequately captured by a continuous function, we also wrote a simple computer programme for a Monte Carlo simulation of the first 12 cell divisions. As expected, this gave almost identical average values to the mathematical model (Fig 7a red lines). However, the distribution of cell numbers at the MBT showed an interesting periodicity in the computer model. Figure 7b shows an example of this periodicity when the error rate was set to 0.002, where distinct peaks of embryos with 0, 2048, 3072 and 3584 cells were evident. We examined the history of the simulated embryos in these peaks and confirmed that they corresponded to embryos where failure had occurred in the first, second, third and fourth cell cycles respectively. The leftward spread of the peaks in Fig 7b represents embryos that subsequently experienced additional cell cycle failures.

**Figure 7.**
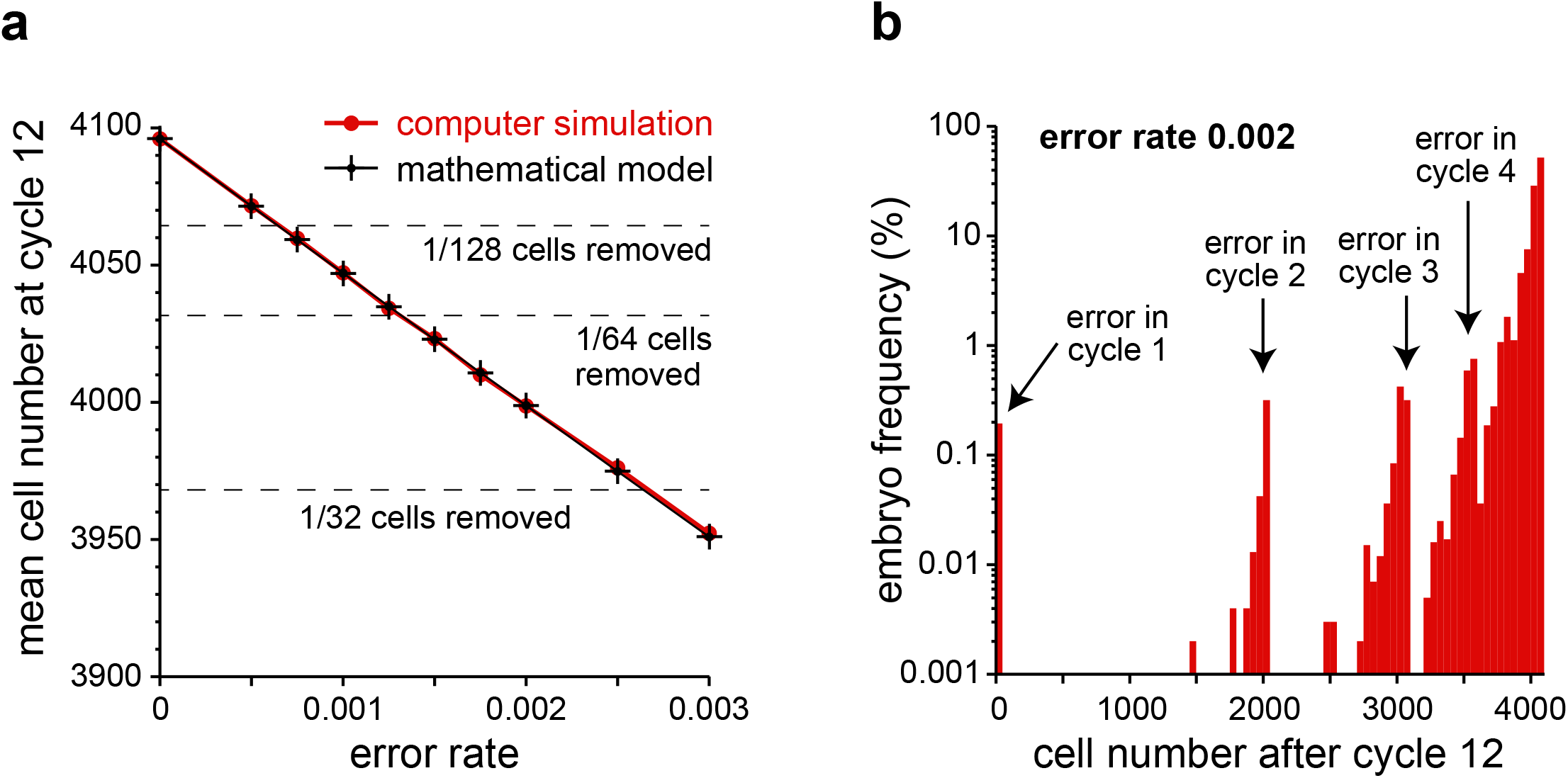
Modelling of embryo survival following loss of cell-lineages. The number of surviving cells (blastomeres) following a series of consecutive cell divisions with a constant probability of cell death per cell division was modelled by a simple mathematical formula (provided in the main text) or a computer programme. **A**. The expected number of surviving cells after 12 cycles of cell division from the mathematical formula (black crosses) or the average of 100,000 iterations of the computer model (red circles). The pre-cell-cycle loss rate is shown on the x axis and the number of cells after cycle 12 is shown on the y axis. Dashed horizontal lines show the values if the maximum number of cells (4,096) is reduced by 1/128, 1/64 or 1/32. **B**. The distribution of surviving cell numbers after 12 rounds of division after 100,000 iterations of the computer model with an error rate of 0.002.

With this estimate for the minimum permissible failure rate for cell division during the cleavage divisions, we determined whether the ‘random completion problem’ (Fig 1b) and the potential danger of double fork stalls (Fig 1a) can be solved by the observed high density of MCM2-7 on chromatin.

The ‘random completion problem’ arises because in early Xenopus embryos, replication initiates randomly with respect to DNA sequence and there is a fixed cell cycle duration that is insensitive to incomplete replication^18,25,33^. This means that if licensed replication origins are more than ∼30 kb apart there will not be enough time for replication to complete before chromosomes are pulled apart at anaphase (Fig 1b). If we assume that MCM2-7 double hexamers are loaded onto inter-nucleosomal linkers, then an equation can be derived for the probability of there being one or more gaps greater than size *N*, where *G* is the genome size (6 Mbp), *k* is the average nucleosomal spacing (200 bp) and is the probability of a linker being licensed by loading MCM2-7 (see Methods for derivation):

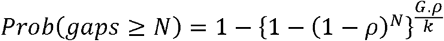

Figure 8a shows this equation plotted for licensing probabilities *ρ* taking values from 0.05 to 0.8. This shows that for linker licensing probabilities greater than ∼0.1, the probability of unlicensed gaps greater than 30 kb is virtually zero. Since we observe that the ratio between MCM2-7 double hexamers and nucleosomes is around 0.8 (Figures 3 and 4) and this *ρ* value is enough to keep the largest unlicensed gaps below 3 kb, we conclude that the density of licensed origins in the early embryo is more than sufficient to solve the ‘random completion problem’.

**Figure 8.**
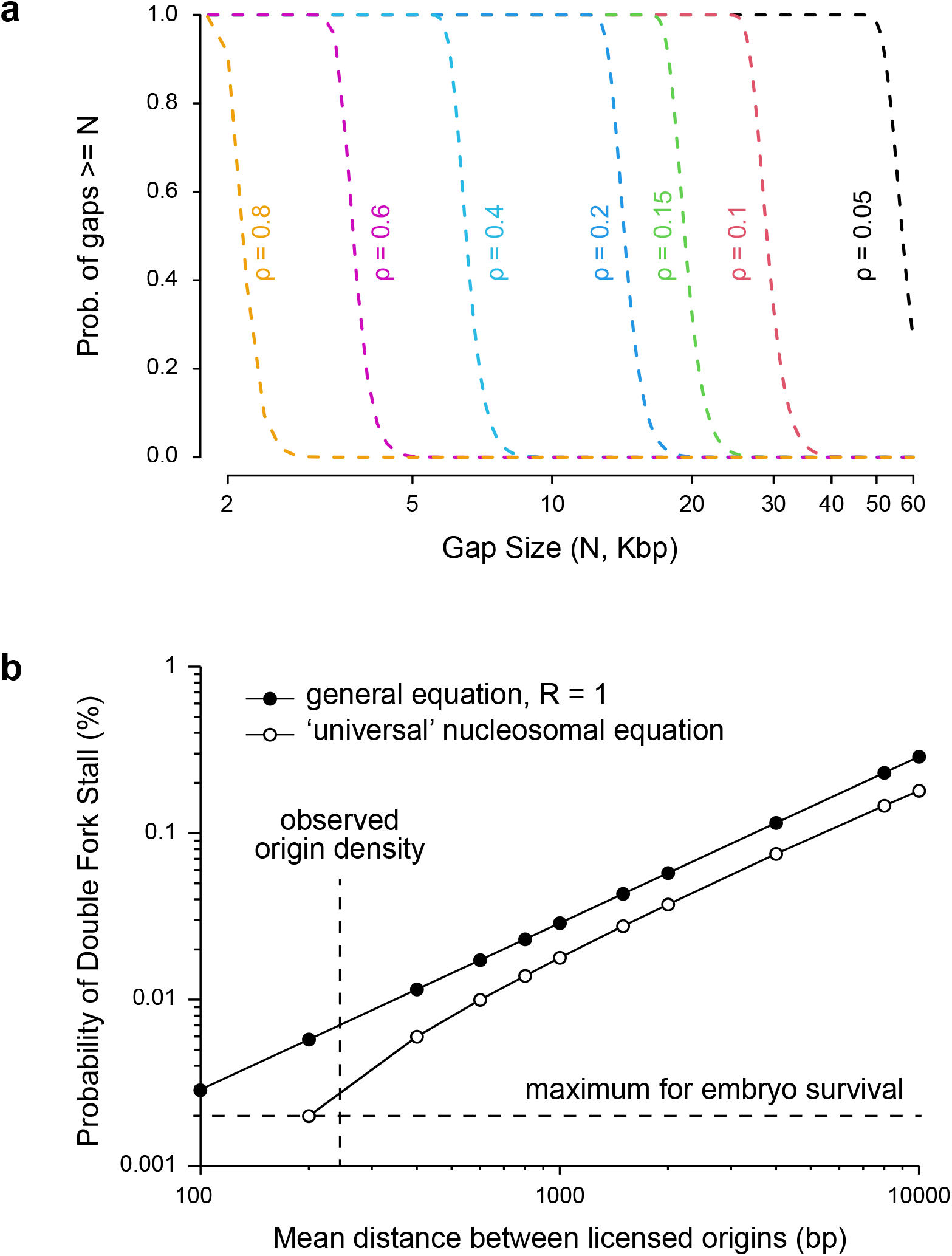
Modelling of unlicensed gaps and double fork stalls. **A**. Mathematical modelling of the largest gap between licensed origins on the *Xenopus* genome, assuming that MCM2-7 double hexamers are loaded onto inter-nucleosomal linker DNA. The x-axis shows unlicensed ‘gaps’ of different sizes; the y-axis shows the probability that a gap of that size occurs. Different lines show different probabilities (ρ) that any given inter-nucleosomal linker is licensed as an origin. **B**. Mathematical modelling of the probability of double fork stalls. The x-axis shows the mean distance between licensed origins; experimental results presented in this paper suggest that the observed value is one MCM2-7 double hexamer every ∼230 bp as indicated by the vertical dashed line. The y-axis shows the probability of a double fork stall; experimental results presented in this paper suggest that for efficient embryo survival, this should be a maximum of ∼0.002 as indicated by the horizontal dashed line. Black circles show the general equation ^4^ when origins are distributed at random (R=1). Open circles show the ‘universal’ equation derived assuming that MCM2-7 double hexamers are loaded onto inter-nucleosomal linker DNA ^7^.

We next considered whether the high density of licensed origins in the early embryo reduces the risk of double fork stalls (DFSs) to a level compatible with embryo survival (Fig 1a). We have previously^4^ derived a generic equation for the probability of one or more DFS occurring genome-wide to be:

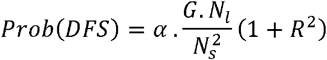

where *α* is a constant 0.24, *G* is the genome size (6 Mbp), *N*_*l*_ is the total number of licensed origins, Ns is the median distance a fork can progress before irreversibly stalling (∼10 Mbp) and *R* is a term that denotes the coefficient of variation of the origin spacing (the ratio of the standard deviation to the mean of the origin separations). Figure 8b (black circles) shows this equation plotted for different densities of licensed origins, assuming that origins are distributed at random (R=1). The observed value of MCM2-7 double hexamer density (∼4 copies per kb, or one double hexamer every 250 bp) gives a predicted DFS probability of 0.007, slightly higher than the maximum tolerated value of ∼0.002 for embryo survival. Note that this equation does not consider the physical footprint of the MCM2-7 double hexamer and allows them to be positioned arbitrarily close to one another.

A better estimate of the distribution of MCM2-7 double hexamers takes into account the presence of nucleosomes on the DNA and the positioning of MCM2-7 in the linker DNA between them. With this assumption we have derived an equation for the probability of one or more DFS occurring genome-wide^7^ to be:

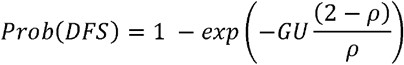

where G is the genome size (6 Mbp), *U* is the ‘universal replication constant’ of ∼3TBp (incorporating an average nucleosome spacing and the mean stall distance) and *ρ* is the probability of a given inter-nucleosome region being occupied by an MCM2-7 double hexamer. Figure 8b (open circles) shows this curve plotted for different densities of licensed origins. At a value for *ρ* of 0.8 (one double hexamer every ∼250 bp) the probability of a DFS is ∼0.003, very close to the maximum tolerated value of ∼0.002 for embryo survival. We therefore propose that the very high density of licensed origins on embryonic chromatin brings the probability of DFSs to values small enough that additional mechanisms for completing DNA synthesis^5,6^ are not required.

The work presented here resolves a number of long-standing questions about DNA replication in the early Xenopus embryo. By using a very high density of origin licensing of almost one per nucleosome, early embryos can reliably complete genome duplication, reducing the probability of DFSs and large inter-origin gaps to levels consistent with embryo survival. Early embryos therefore do not need complex and time-consuming post-S phase or mitotic pathways for completing DNA replication, thus ensuring that they can divide very rapidly. This level of origin licensing does not perturb nucleosome spacing, consistent with MCM2-7 being loaded onto nucleosome linker DNA. The observation that almost any DNA sequence introduced into Xenopus eggs can serve as a functional replication origin is likely to be a by-product of the very high density of MCM2-7 on DNA. However, the extremely high abundance of MCM2-7 on chromatin is likely to present a huge barrier to transcription. We therefore suggest that animal cells with large genomes can take two radically-different approaches to ensuring complete genome duplication: either to have relatively long cell cycles whose stages can be lengthened to deal with replication problems – the situation in the vast majority of somatic cells – or to have short cell cycles but in the absence of transcription – the situation in many rapidly-dividing early embryos.

## Acknowledgements

This work was supported by CR-UK programme grant C303/A14301 and Wellcome Trust Investigator award WT096598MA. We would like to acknowledge help from Dr. Sara ten Have and Kelly Hodge of the GRE Proteomics Support Team at the University of Dundee (Wellcome Trust Strategic Award 097945/Z/11/Z). The authors declare that they have no conflicts of interest with the contents of this article.

## Author contributions

PJG performed all of the egg extract experiments with the exception of quantifying the MCMs in egg extract; JK performed the embryo experiments; MAM performed the mathematical modelling; GK performed the proteomics analysis; KC developed the chromatin isolation procedure together with PJG; AJS performed the MCM quantification and PJG analysed the data; JJB led the project and performed the computer modelling. JJB and PJG prepared the figures and wrote the manuscript.

## Materials and Methods

Requests for resources, reagents and further information should be directed to and will be fulfilled by, the corresponding author, J. Julian Blow.

### Antibodies & reagents

Anti-histone H3 FL-136 rabbit (Santa Cruz sc-10809) was used to determine chromatin recovery from egg extract. Antibodies used in this study raised against *Xenopus* proteins were: MCM3 and MCM6 ^48^; ORC1 ^44^; Cdc6 ^45,69^. Anti-rabbit IgG was from New England Biolabs. Recombinant geminin^DEL^ was produced as described ^70^ and p27^KIP1^ was produced as described in ^71^. *Triticum vulgaris* Wheat Germ Agglutinin was purchased from Sigma-Aldrich (L9640).

### Xenopus laevis

Wild type, sexually mature (≥ 1 year old) female and male South African Clawed frogs (*Xenopus laevis*) born and reared in the UK (University of Portsmouth) were used in this study for the production of unfertilised eggs (from which extracts were prepared) and sperm, respectively. Frogs were maintained at 19°C in particulate filtered, dechlorinated water, at a density of ≤15 animals per 60 l tank, in a purpose built ‘aquacentre’ and were maintained by professional staff at the University of Dundee adhering to Home Office (UK Government) animal husbandry guidelines; the animals have access to a Home Office (UK Government) approved veterinary surgeon. The frogs were fed a vegetable and fish based diet (Aquatic Diets 3, Mazuri Zoo Foods) 2-3 times per week, as required.

### Xenopus egg extract methods

Metaphase-arrested *Xenopus laevis* egg extract and demembranated *Xenopus* sperm nuclei were prepared as described^23,72^. Female frogs were primed with 50 units of Folligon (Pregnant Mare Serum Gonadotrophin) 3 days before the eggs were required to increase the number of stage 6 mature oocytes and 2 days later, were injected with 500 units Chorulon (Chorionic Gonadotrophin, Intervet) to induce ovulation. Frogs were placed in individual laying tanks at 18-21°C in 2 l 1x MMR egg laying buffer, prepared from a 10x stock (1 M NaCl, 20 mM KCl, 10 mM MgCl_2_, 20 mM CaCl_2_,1 mM EDTA, 50 mM HEPES-NaOH, pH 7.8). The following morning, eggs were collected and rinsed in 1x MMR to remove any non-egg debris. Washed eggs were dejellied in 2% w/v cysteine (pH 7.8), washed in XBE2 (1x XB salts, 1.71% w:v sucrose, 5 mM K-EGTA, 10 mM HEPES-KOH, pH 7.7; 10x XB salts: 2 M KCl, 40 mM MgCl_2_, 2 mM CaCl_2_) and then into XBE2 containing 10 μg/ml leupeptin, pepstatin and aprotinin. Dejellied and washed eggs were centrifuged in 14 ml tubes, containing 1 ml XBE2 plus protease inhibitors containing 100 μg/ml cytochalasin D, at 1400 x *g* in a swinging bucket rotor for 1 min at 16°C to pack the eggs, after which excess buffer and dead eggs were removed. Packed eggs were crushed by centrifugation at 16,000 x *g* in a swinging bucket rotor for 10 min at 16°C. The dirty brown cytoplasmic layer was collected using a 20G needle and a 1 ml syringe via side puncture. From this point onwards the extract was kept on ice. The crude extract was supplemented with cytochalasin D, leupeptin, pepstatin and aprotinin all to a final concentration of 10 μg/ml, 1:80 dilution of Energy Regenerator (1 M phosphocreatine disodium salt, 600 μg/ml creatine phosphokinase in 10 mM HEPES-KOH pH 7.6) and 15% v:v LFB1/50 (10% w:v sucrose, 50 mM KCl, 2 mM MgCl_2_, 1 mM EGTA, 2 mM DTT, 20 mM K_2_HPO4/KH_2_PO4 pH 8.0, 40 mM HEPES-KOH, pH 8.0). The extract was clarified by centrifugation at 84,000 x *g* in a pre-cooled SW55 rotor swinging bucket rotor at 4°C for 20 min. The golden cytoplasmic layer was recovered, supplemented with glycerol to 2% v/v and frozen in aliquots in liquid nitrogen and stored at -80°C until required.

Sperm was recovered from testes isolated from male frogs post mortem following a lethal dose of anaesthetic (0.2% w:v Tricaine mesylate MS222, ∼0.5% w:v NaHCO3, to pH 7.5). Isolated testes were washed carefully to avoid bursting in EB (50 mM KCl, 5 mM MgCl_2_, 2 mM dithiothreitol or β-mercaptoethanol, 50 mM HEPES-KOH, pH 7.6), prior to being finely chopped with a clean razor blade in fresh EB. Recovered lysate was filtered through a 25 μm nylon membrane to remove particulate matter. Filtered sperm was centrifuged at 2,000 x *g* at 4°C for 5 min; selective resuspension of the sperm pellet allowed separation of the sperm from contaminating erythrocytes; the resuspended sperm was respun and the pellet resuspended in 0.5 ml SuNaSp (0.25 M sucrose, 75 mM NaCl, 0.5 mM spermidine, 0.15 mM spermine, 15 mM HEPES-KOH, pH 7.6) per testis. The sperm was demembranated with the addition of 25 μl per testis lysolecithin (5 mg/ml, in H_2_O) for 10 min at room temperature. Demembranated sperm were respun and resuspended in SuNaSp plus 3% w/v BSA to quench the demembranation reaction. Quenched sperm were respun and resuspended in 100 μl EB plus 30% glycerol per testis, counted using a haemocytometer and stored at -80°C.

Extracts were supplemented with 250 μg/ml cycloheximide, 25 mM phosphocreatine and 15 μg/ml creatine phosphokinase and incubated with 0.3 mM CaCl_2_ for 15 min to trigger release from metaphase arrest. Desalted egg extract was prepared as described ^57^.

### Chromatin Isolation from extract

Chromatin isolations from *Xenopus* egg extract was undertaken using low adhesion Eppendorf tubes, as described ^23,72^: reactions were stopped by the addition of 400 μl of ice-cold NIBTX (50 mM KCl, 50 mM Hepes KOH pH 7.6, 5 mM MgCl_2_, 2 mM DDT, 0.5 mM spermidine 3HCl, 0.15 mM spermine 4HCl, 0.1% Triton X-100). This was underlayered with 100 μl 15% sucrose in NIBTX. The tubes were spun at 2100 x *g* for 5 min at 4°C in a swinging bucket rotor. The buffer above the sucrose cushion was removed and the surface of the cushion washed with 200 μl NIBTX before removing the cushion down to ∼15 μl. The tubes were then spun at 13000 x *g* for 2 min in a fixed angle rotor to focus the chromatin pellet and following this, all the buffer was removed. The chromatin pellet was then resuspended in Laemmli loading buffer and was subjected to SDS-PAGE using 4–12% Bis-Tris gradient SDS–PAGE gel (Invitrogen). Control experiments determined that this protocol consistently recovered ∼90% of chromatin added to the extract (data not shown).

### Xenopus laevis embryo experiments

Highly synchronized embryos of *Xenopus laevis* (Linnaeus) were obtained by *in vitro* fertilization according to ^73^. Fertilization rate was >95%. To remove the viteline layer embryos were placed in 2% cysteine in 0.1xMMR (10x stock: 20 mM CaCl_2_, 50 mM HEPES, 20 mM KCl, 10 mM MgCl_2_, 1 M NaCl, pH 7.8) and washed with 1x MMR and then 0.1x MMR. Further development continued in fresh 0.1x MMR.

DNAse (Roche) stock was made in the buffer containing 400 mM TRIS-HCl, 100 mM NaCl, 60 mM MgCl_2_ and 10 mM CaCl_2_, pH 7.9. DNAse was then diluted in an injection buffer (25 mM TRIS-HCl, 50% glycerol v/v, pH 7.6). 30 nl, containing 1U of DNAse, was injected into a single cell of 2-cell (stage 2), 4-cell (stage 3), 8-cell (stage 4), 16-cell (stage 5), 32-cell (stage 6), 64-cell (stage 6.5, MBT) or 128-cell (stage 7, post MBT) embryo. The non-injected part of an embryo served as a first control and the development of non-injected embryos served as a control from the same batch of embryos. A small number of embryos were injected with the same buffer but without DNAse. For practical reasons this control was done from the next fertilized batch of embryos. No significant difference to the non-injected embryos was observed, however 1 of 5 injected embryos died soon after injection in stage 3 and 4.

Development was followed by live imaging using a Motic stereomicroscope. One of the embryos from each injected group was fixed at MBT, 24 hrs and together with all that survived at 50 hrs of development. MEMFA salts (10x stock: 1M MOPS, 20 mM EGTA, 10mM MgSO_4_, pH 7.4) and 37% formaldehyde followed by two washings in methanol ^74^ was used in this procedure. Fixed embryos were further imaged using Leica fluorescence Macroimaging system M205FA with digital high sensitivity DFC310FX cooled camera and LasA program.

### Immunoblotting & Gel Staining

Samples were prepared in 6x Laemmli sample buffer, heated at 99°C for 4 min, and separated on a 4-12% BisTris NuPage gels (Invitrogen) in 1x MOPS running buffer (Invitrogen). Proteins were transferred onto PVDF membrane (Cytiva) and detected using either enhanced chemiluminescence detection (SuperSignal West Pico Chemiluminescent Substrate; ThermoFisher) or fluorescent secondary antibody (ThermoFisher); images were captured using either an ImageQuant LAS4000 (Amersham (Cytiva)) CCD - for chemiluminescence - or an Odyssey CLx (LiCor) - for fluorescent secondary antibodies.

For protein visualisation gels were stained with either SYPRO Ruby (Thermo Fisher) and de-stained with 10% ethanol, 7% acetic acid, in H_2_O or with Coomassie R-250 and de-stained with 40% ethanol, 10% acetic acid in H_2_O. SYPRO Ruby stained gel images were captured using a Typhoon laser-scanning gel-imaging system (Cytiva) and quantified using Image Studio Lite (LiCor).

Immunoblots were quantified using Image Studio Lite (LiCor) and Coommassie gels were quantified using GelEval (FrogDance Software).

### Orc1 and Cdc6 chromatin saturation

Egg extract was first treated with apyrase to remove ATP. Metaphase-arrested extract was supplemented with cycloheximide only (no Energy Regenerator was added) and released into interphase for 5 min by the addition of CaCl_2_, as described above. The extract was then supplemented with 0.01 U ml^-1^ of Apyrase (Sigma A6410) for 10 min, followed by the addition of 2.5 mM ATP-γ-S. Different volumes of ATP-γ-S supplemented extract were added to 200 ng of sperm nuclei and incubated for 20 min. Chromatin was isolated, as described above. Recovered samples were resuspended in 6x Laemmli loading buffer and were run on an SDS-PAGE gel, together with 0.5 μl of whole egg extract to serve as a control for quantification and then immunoblotted and quantified as described above. The lower portion of the gel was stained with SYPRO Ruby to detect histones as a control for chromatin recovery.

### MCM3 and MCM6 chromatin saturation - immunoblotting

Metaphase-arrested egg extract was supplemented with cycloheximide and Energy Regenerator and was activated to enter interphase with CaCl_2_, as described above. The activated extract was supplemented with 2 mg ml^-1^ wheat germ agglutinin and 10 ng μl^-1^ sperm nuclei; at the indicated times the licensing reaction was stopped by the addition of 100 nM geminin^DEL^. All samples were isolated, as described above, following completion of the 20 minute time course. Recovered samples were resuspended in 6x Laemmli loading buffer and 30 ng of chromatin from each sample was run on an SDS-PAGE gel, together with 25 nl of whole egg extract to serve as a control for quantification and then immunoblotted and quantified as described above. Chromatin recovery was visualised by immunoblotting for Histone H3.

### MCM2 and MCM5 chromatin saturation - SYPRO Ruby

The *Xenopus* haploid genome corresponds to 3.15 pg DNA. Following fertilisation and the completion of eleven of the twelve cleavage cell divisions the egg contains ∼12.9 ng of DNA. Since each Xenopus egg yields ∼0.5 μl of soluble material the DNA concentration in the egg for the final S phase prior to MBT is ∼26 ng μl^-1^.

Metaphase-arrested egg extract was supplemented with cycloheximide and Energy Regenerator and was activated to enter interphase with CaCl_2_, as described above. The activated extract was supplemented, or not, with 26 ng μl^-1^ sperm nuclei ±100 nM geminin^DEL^ for 20 min. Samples were isolated, as described above except that samples were resuspended in 70 mM KCl and a 20% sucrose cushion was used. Recovered samples were resuspended in 6x Laemmli loading buffer and 50 ng of chromatin from each sample was run on an SDS-PAGE gel, proteins were visualised using SYPRO Ruby and gel images were captured and quantified, as described above.

To enable quantification of MCM2 and MCM5 on chromatin following one round of genome duplication, sperm chromatin was recovered following the completion of S phase and extract conversion into metaphase. Metaphase-arrested egg extract was supplemented with cycloheximide and ER and was activated to enter interphase with CaCl_2_, as described above. Following release from metaphase, interphase extract was supplemented with 10 ng μl^-1^ sperm nuclei and incubated for 2 hours to allow complete genome duplication. The replicated chromatin was converted into condensed metaphase chromosomes following the addition of 2 volumes of metaphase-arrested extract and a 90 min incubation. Chromatin was isolated by diluting the reaction with 1 ml NIB50 (50 mM KCl, 50 mM Hepes KOH pH 7.6, 5 mM MgCl_2_, 2 mM DDT, 0.5 mM spermidine 3HCl, 0.15 mM spermine 4HCl), underlaid with 100 μl NIB50 containing 15% sucrose, underlaid with 5 μl NIB50 containing 30% glycerol and spun at 6000 x g, 5 minutes at 4°C in a swinging bucket rotor bench centrifuge. The buffer above the sucrose cushion was washed with 200 μl NIB50 and the buffer was removed leaving ∼15 μl sample. Chromatin was gently re-suspended by inversion and was immediately added to a second interphase extract and incubated for 20 min, re-isolated and run on an SDS-PAGE gel, visualised with SYPRO Ruby and MCM2 and MCM5 quantified, as described above. Chromatin recovered for use in a second extract incubation is isolated in the total absence of detergent.

### Micrococcal Nuclease (MNase) digestion of chromatin

Metaphase-arrested egg extract was supplemented with cycloheximide and Energy Regenerator and was activated to enter interphase with CaCl_2_, as described above. Interphase extract was supplemented with 2 mg ml-1 wheat germ agglutinin and 10 ng μl^-1^ sperm nuclei ±100 nM geminin^DEL^ for 20 min or 10 ng μl^-1^ sperm chromatin alone for 2 hours to allow complete genome duplication.

50 μl egg extract per digestion timepoint per condition was diluted 10 fold in room temperature buffer XN (50 mM HEPES-KOH, pH 7.0; 250 mM sucrose, 75 nM NaCl, 0.5 mM spermidine 3 HCl and 0.15 mM spermine 4 HCl), underlayered with 5 μl 30% glycerol in buffer XN and chromatin recovered, in the absence of a sucrose cushion, by centrifugation at 4000 x *g* for 5 min in a swinging bucket rotor. After removing the supernatant, each pellet was gently resuspended into 50 μl MNase buffer (10 mM TrisCl, pH 7.5; 80 mM NaCl; 2.0 mM CaCl2, 25 % v/v glycerol) and supplemented with 1 U MNase ng^-1^ DNA and incubated for the indicated time. Digestion was stopped with the addition of 350 μl MNase STOP buffer (0.1 M TrisCl, pH 8.0; 5 mM EDTA; 0.2 M NaCl, 0.2 % w/v SDS and 0.5 mg ml^-1^ proteinase K) and incubated for 2 hr at 37°C. 200 μl 4.5 M NaCl was then added and samples mixed on a rotating wheel for 10 min before being centrifuged at 2,000 x *g* in a microcentrifuge. The recovered supernatant was removed to a 2 ml tube and 1200 μl 100% ethanol was added, the sample mixed and incubated overnight at -20°C to facilitate DNA precipitation. Precipitated DNA was recovered by centrifugation at 5,000 x *g* in a microfuge, the pellet was washed with 70 % ethanol, spun and then dried for 15 min at room temperature in a fume hood. The dried pellet was resuspended in 20 μl TE supplemented with 0.1 mg ml-1 RNase A and incubated for 2 hours at room temperature before being run on a 1.0 % TAE gel together with a 100 bp ladder (NEB) for ∼90 min at 100 V. The gel was stained with Ethidium Bromide and visualised using a Smart 3 GelDoc System (VWR).

### Proteomics

Metaphase-arrested egg extract was supplemented with cycloheximide and Energy Regenerator and was activated to enter interphase with CaCl2, as described above. The activated extract was supplemented with 10 ng μl^-1^ sperm nuclei, ±100 nM geminin^DEL^, ±100 nM p27^KIP1^ and incubated for 2 hours to allow complete genome duplication. Samples were prepared in triplicate and analysed with respect to starting extract volume not final DNA concentration. Mass spectrometry was performed as described^66^ .

### Mathematical Modelling

For deriving the probability of having one or more gaps bigger than a given size of *N* bp, we define *ρ* as the probability of a nucleosome being licensed with MCM2-7, *G* as the genome size and *k* as the average nucleosome spacing in the genome. The expected number of nucleosomes with MCM2-7 is 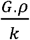. So, the probability of having a gap of *n* unlicensed nucleosomes, *p*_*n*_, is given by

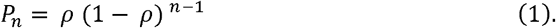

Let 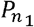 and 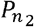 be the probabilities that the first and second gap on the genome has *n*_1_ and *n*_2_ numbers of unlicensed nucleosomes, and so on. Then, the probability that the <u>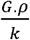</u> gaps have sizes 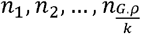 is given by

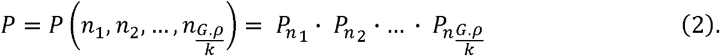

Now, the probability that a single independent gap *n* has size smaller than a particular size *N* is given by

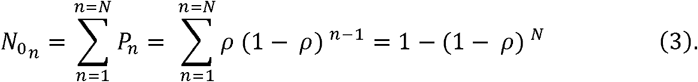

The probability that all the gaps 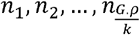 are smaller than *N* is the product of all independent probabilities for each gap being less than size *N* that are given by 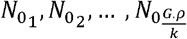. Hence, the probability for all gaps genome-wide are less than size *N* is

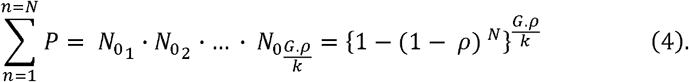

So the probability of one or more gaps to be bigger than size *N* is given by

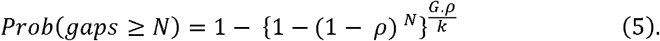

### Computer Modelling

The computer programme for a Monte Carlo simulation of the first 12 cell divisions was written in Swift 5.0 using the Xcode development environment. The simulation function code is provided in Supplementary Figure S6.

