## Supplemental figure 1-6 for "High levels of origin licensing during *Xenopus* cleavage divisions ensures complete and timely genome duplication"

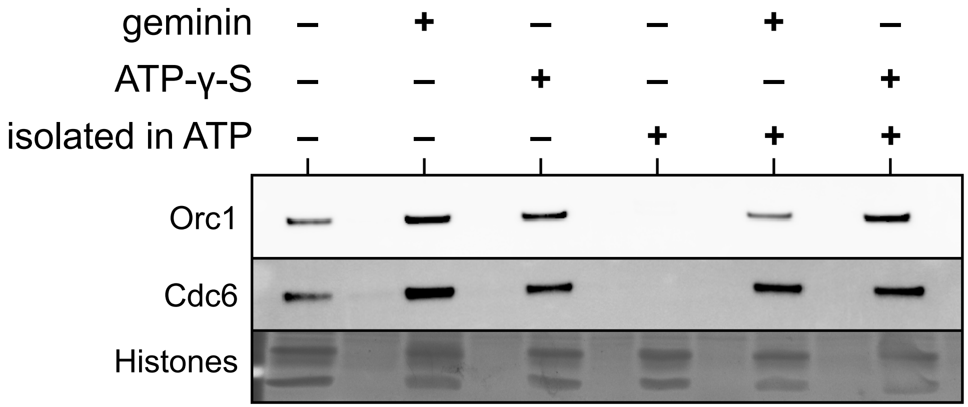


**Supplementary Figure S1.**
Interphase Xenopus egg extract was either supplemented or not with 100 nM gemininDEL or treated with Apyrase to remove ATP (Gardner, et al) and then supplemented with 2.5 mM Mg•ATP-γ-S; Xenopus laevis sperm chromatin was added to the treated extract and incubated for 20 mins before isolation over a sucrose cushion in either the presence of absence of 2.5 mM Mg•ATP. Isolated chromatin was subjected to SDS-PAGE. The upper portion of the gel was immunoblotted and probed with antibodies raised against Orc1 and Cdc6 and the lower portion of the gel was stained with Coomassie Blue to visualise histones which serve as a control for

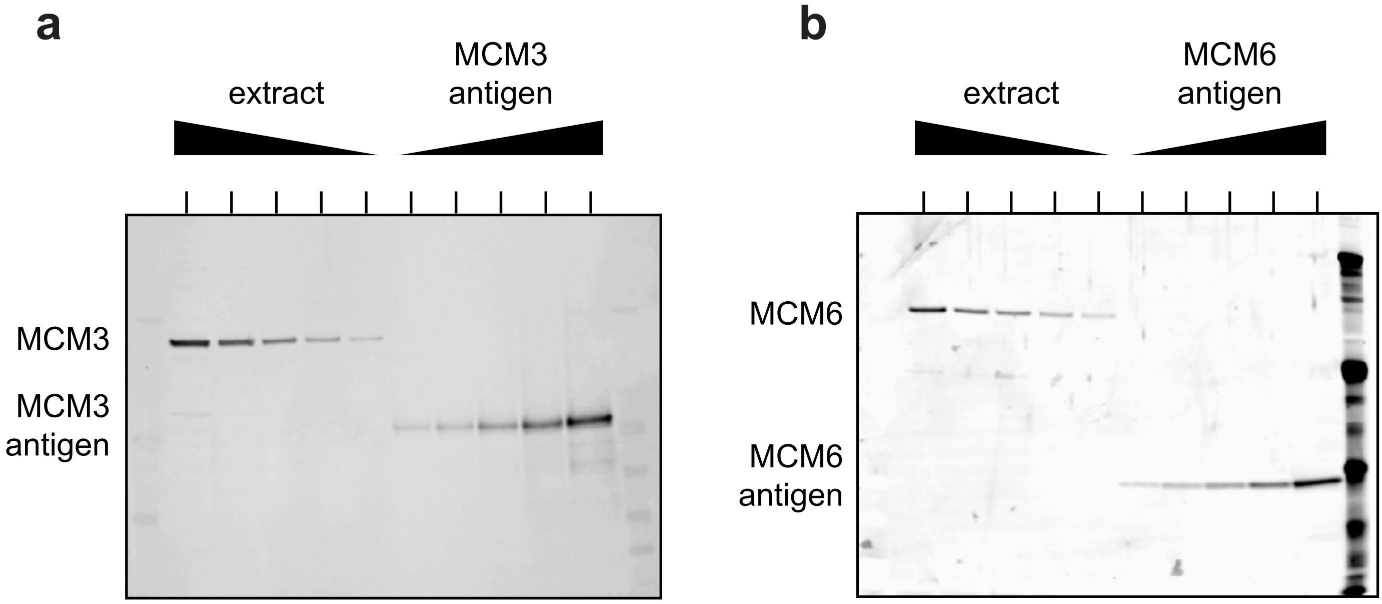


c

| | [antigen] (μM) | [extract] / [antigen] | [Mcm] (μM) |
|---|---|---|---|
| MCM3 | 12.6 | 1.48 ± 0.20 | 18.6 ± 2.5 |
| MCM6 | 27.7 | 0.585 ± 0.074 | 16.2 ± 2.1 |

**Supplementary Figure S2.**
**a.** Exemplar 2-fold dilutions of Xenopus egg extract (lowest dilution 1/8) and MCM3 antigen (lowest dilution 1/3) immunblotted with antibodies against MCM3.
**b.** Exemplar 2-fold dilutions of Xenopus egg extract (lowest dilution 1/32) and MCM6 antigen (lowest dilution 1/32) immunblotted with antibodies against MCM6.
**c.** The signals were quantified in 4 separate experiments to give a ratio between the concentration of the antigen and the concentration of the protein in extract, allowing the concentration of protein in extract to be estimated. Vlaues are shown ± standard deviation.

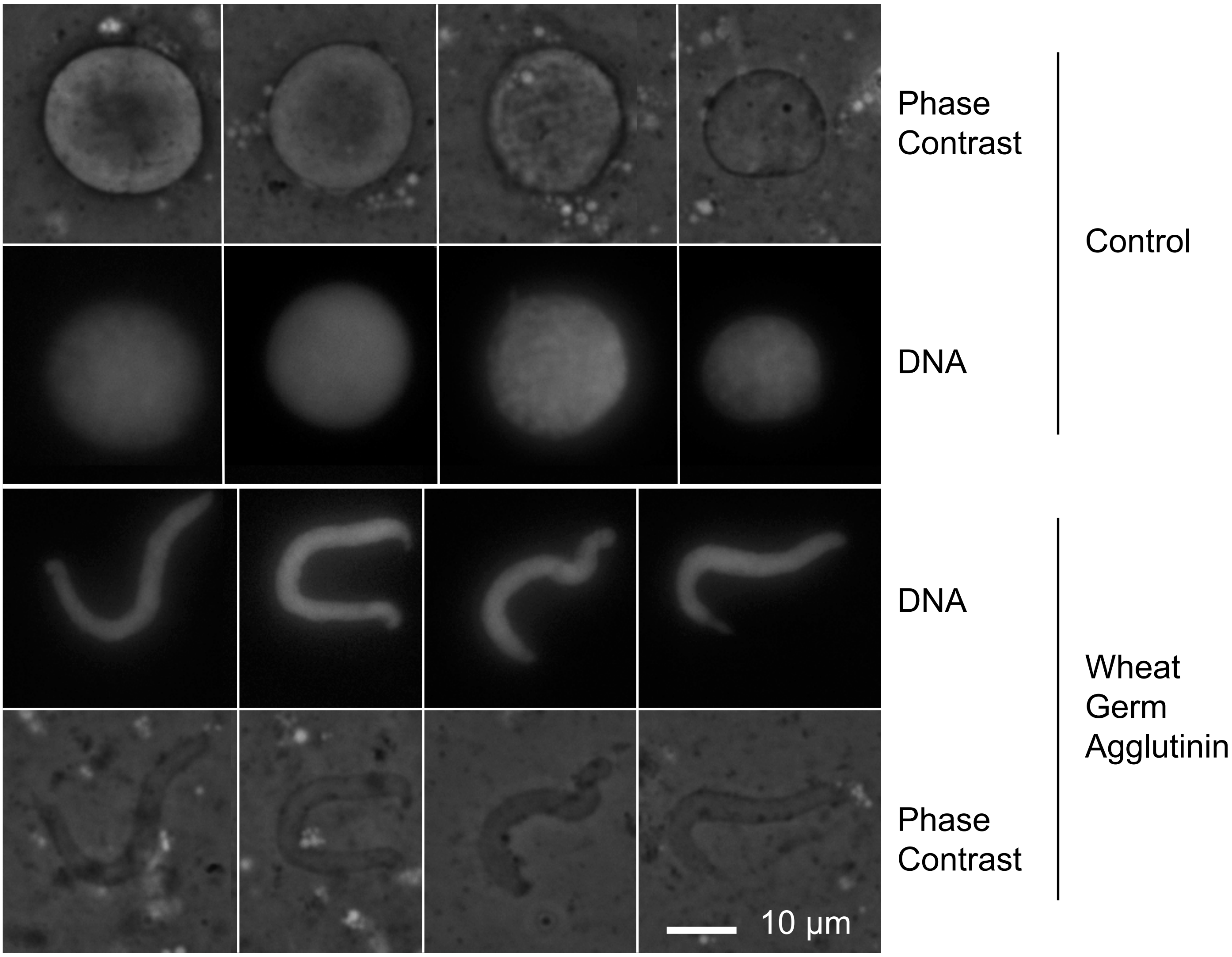


**Supplementary Figure S3.**
Interphase egg extract was supplemented or not with 2 mg $ml^{-1}$ wheat germ agglutinin and then Xenopus sperm nuclei were added to a final concentration of 10 ng $\mu l^{-1}$ and extracts were incubated for 20 mins; the state of nuclear formation was determined by phase contrast microscopy and DNA visualised following Hoescht 33342 staining by fluorescence microscopy. 4 exemplar nuclei are shown for each condition. The scale bar represents 10 µm.

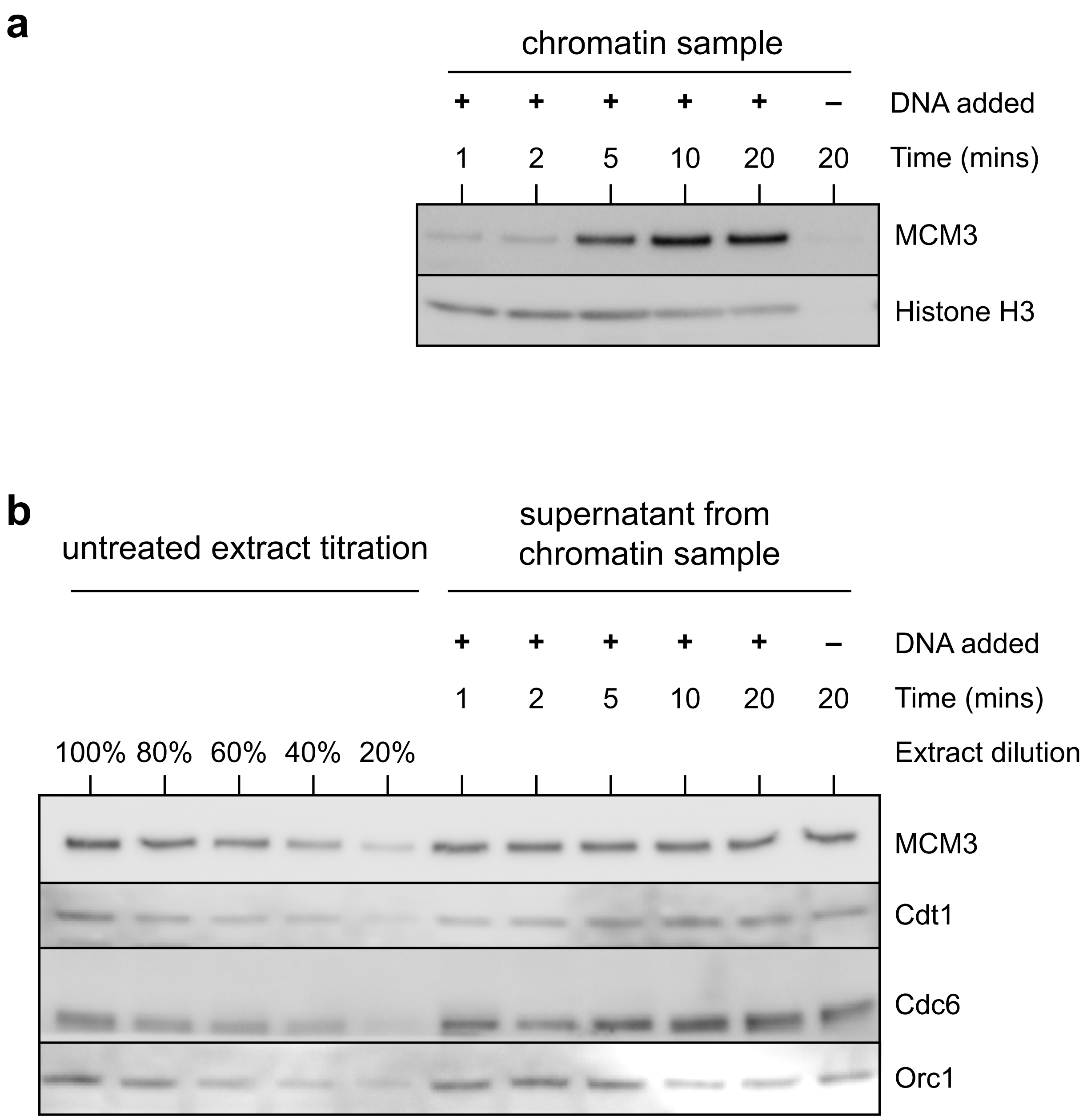


**Supplementary Figure S4.**
Interphase egg extract was incubated for 1, 2, 5, 10 or 20 mins plus or minus Xenopus sperm nuclei at a final DNA concentration of 10 ng µl$^{-1}$. Chromatin was isolated from the extract by centrifugation. **a)** Chromatin was immunblotted for Mcm3 and histone H3. **b)** The supernatant was loaded onto a gel and immunblotted for Orc1, Cdc6, Cdt1 and Mcm3. As a control, an equal volume of untreated extract titrated to 100, 80, 60, 40 or 20% was also loaded on the gel.

!"#$%&()*+,-*./*0!"1 *

**a**

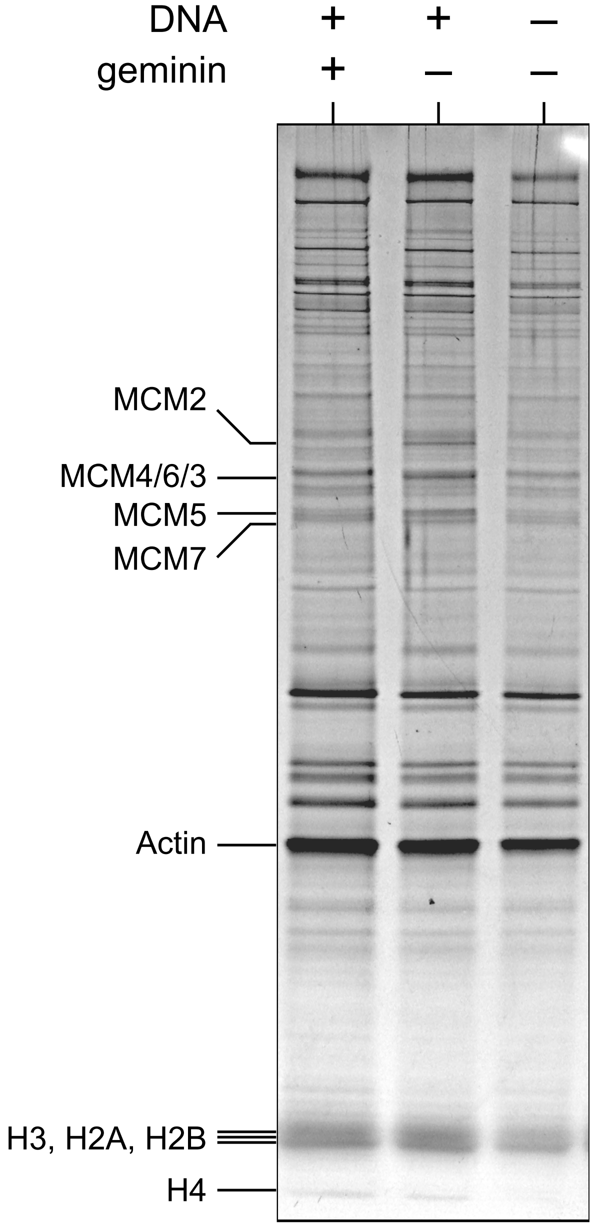


**b**

| | band intensity | n = 13 K+R+H | protein quantity |
|---|---|---|---|
| **MCM2** | 4.41 ± 0.56 | 137 | 0.869 ± 0.110 |
| **MCM5** | 3.23 ± 0.65 | 121 | 0.721 ± 0.145 |
| **H4** | 1.0 ± 0.0 | 27 | 1.0 ± 0.0 |

**Supplementary Figure S5.**
Chromatin was added to an interphase egg extract to a final concentration of 10 ng μl$^{-1}$ and incubated for 2 hours to allow complete genome duplication. 2 volumes of metaphase extract was added and incubated for 90 mins to induce nuclear envelope breakdown and the formation of metaphase chromosomes; chromosomes were isolated over a sucrose cushion onto a glycerol cushion. The recovered chromatin was added to a second interphase egg extract in the presence or absence of 100 nM geminin$^{DEL}$ and incubated for 20 mins. Chromatin was isolated over a sucrose cushion and subjected to SDS-PAGE; gels were stained with SYPRO Ruby to visualise recovered proteins. The position of Mcm2-7 and the histones is indicated

```
func performSimulation(cellCycles: Int, errorRate: Double, simNumber: Int) -> ([Int],[Int],[Int]) {
  /*   Inputs to the function:
       cellCycles: the number of embryonic cell cycles in the simulation
       errorRate: the per-cell-cycle probability of failing to pass through the cell cycle; if an error occurs,
            the cell dies
       simNumber: the number of simulations to be performed

       Outputs from the function:
       embryoCellCounts: array of the count of embryos having a cell number as given by array index
       embryosWithDefectsAtCycle:  the count of embryos experiencing errors in the cell cycle as given
            by array index
       totalCellCountsForEmbryosWithDefectsAtCycle: the total count of cells in embryos experiencing
            errors in the cell cycle as given by array index      */

  var cellNumber = 1
  for _ in 0 ..< cellCycles { cellNumber *= 2 }
  let maxCellNumber = cellNumber                    // the final number of cells in each embryo if no errors
  var embryoCellCounts = [Int](repeating: 0, count: maxCellNumber+1)      // initialise output arrays
  var embryosWithDefectsAtCycle = [Int](repeating: 0, count: cellCycles)
  var totalCellCountsForEmbryosWithDefectsAtCycle = [Int](repeating: 0, count: cellCycles)

  for _ in 0 ..< simNumber {                    // loop through each simulation
    var cellNumber = 1                          // start with 1 cell in the embryo
    var defectOccurredAtCycle = [Bool](repeating: false, count: cellCycles)         // record of errors
    for cellCycleIndex in 0 ..< cellCycles {        // loop through each cell cycle, starts at 0
      var newCellNumber = 0
      for _ in 0 ..< cellNumber {               // attempt to divide each cell
        if Double.random(in: 0...1) > errorRate { newCellNumber += 2 }       // if successful division
        else { defectOccurredAtCycle[cellCycleIndex] = true }
      }
      cellNumber = newCellNumber
    }

    embryoCellCounts[cellNumber] += 1              // record the cell number in the frequency array

    for cellCycleIndex in 0 ..< cellCycles {             // record where the cell cycle failures were and link to
                                                            final cell number in embryo
      if defectOccurredAtCycle[cellCycleIndex] {
        embryosWithDefectsAtCycle[cellCycleIndex] += 1
        totalCellCountsForEmbryosWithDefectsAtCycle[cellCycleIndex] += cellNumber
      }
    }
  }
  return
(embryoCellCounts, embryosWithDefectsAtCycle, totalCellCountsForEmbryosWithDefectsAtCycle)
}
```
